# A constitutive model for the remodelling erythrocyte membrane skeleton during the active invasion by the malaria merozoite

**DOI:** 10.1101/2023.06.13.544728

**Authors:** Chimwemwe Msosa, Tamer Abdalrahman, Thomas Franz

## Abstract

Malaria merozoites phosphorylate erythrocyte membrane proteins to breach the membrane during invasion. This study aimed to develop a constitutive model for erythrocyte membrane phosphorylation that reduces the membrane’s elastic modulus and resistance to merozoite invasion. The hyperelastic Mooney Rivlin constitutive model was adapted by adding an exponential term to represent the mechanical effect of erythrocyte membrane phosphorylation. The modified algorithm was verified with the unmodified Mooney Rivlin model for the intact erythrocyte membrane and used to predict erythrocyte membrane stress for equi-biaxial membrane strain up to 1.1 for different severity of phosphorylation damage. The stability of the damage model was assessed using the Drucker criterion for equi-biaxial strain up to 2.0. Strain and stress predicted with the developed damage model and the Mooney Rivlin model agreed for the intact erythrocyte membrane. The membrane stress at a strain of 1.1 decreased by 42% for minor and 95% for severe erythrocyte membrane damage. The stability strain threshold of the damage model was 1.98 for minor and 1.19 for severe membrane damage. The developed model can represent different degrees of erythrocyte membrane damage through phosphorylation by a malaria merozoite. The model will enable *in silico* investigations of the invasiveness of malaria merozoites.

## 1 Introduction

Like in many biological cells, the erythrocyte membrane acts as a mechanical structure and chemical barrier, which allows the erythrocyte to undergo large nonlinear elastic deformations and regulate transport processes across it. Hence, any persistent disruption of the erythrocyte membrane can impair various physiological functions of the erythrocyte.

The erythrocyte membrane skeleton is a dense network of proteins. This network provides the erythrocyte membrane’s mechanical strength, allowing the cell to withstand mechanical loads as it flows through the narrow blood vessels [1-3]. It comprises a unique arrangement of spectrin, F-actin, protein 4.1, and ankyrin with direct and indirect connections to the lipid bilayer [4]. Spectrin proteins form a large part of the erythrocyte membrane skeleton and are responsible for the elastic deformation of the cell [5]. Apart from providing mechanical strength to the erythrocyte membrane, the spectrin network is the structural platform for stabilising and activating membrane channels, receptors, and transporters [6]. The importance of the protein network of the erythrocyte membrane skeleton in maintaining the discoid shape and elasticity of a healthy erythrocyte has been highlighted in various studies involving erythrocytes with hereditary disorders [7, 8].

Malaria merozoites evade its destruction from the host immune system by invading a human erythrocyte [9]. The merozoite must invade the human erythrocyte without breaching the membrane barrier to avoid erythrocyte cell death. Erythrocyte membrane damage can be categorised into two types: (1) biochemical damage involving a chemically induced kinetic reaction, leading to the reorganisation of the erythrocyte membrane skeleton, and (2) biomechanical damage due to excessive membrane deformation.

During the merozoite invasion, the erythrocyte membrane’s chemical damage is induced by the erythrocyte binding antigen (EBA-175) binding to the glycophorin A (GPA) erythrocyte receptor that initiates biochemical signalling cascades. The complex signalling cascade involves the phosphorylation of critical proteins of the membrane skeleton, such as ankyrin and spectrin, resulting in considerable changes in the mechanical properties of the erythrocyte membrane. EBA-175 is a plasmodium falciparum merozoite antigen secreted by micronemes that binds to the GPA through the apical region of the merozoite. It has been demonstrated by nano-indentation that EBA-175 reduces the elastic modulus and thus increases the deformability of the erythrocyte membrane [10]. The primary mechanism associated with the reduction in the erythrocyte membrane elastic modulus is EBA-175-activated phosphorylation. In general, phosphorylation and dephosphorylation of membrane proteins is an essential mechanism by which erythrocyte membrane properties are regulated [11].

Previous studies have described molecular factors altering the elastic behaviour of the erythrocyte membrane [10, 12]. However, the impact of the erythrocyte membrane remodelling [13] on the merozoite’s invasiveness is not well understood. Theoretical and computational models allow the investigation of complex biological systems and complement experimental studies [14, 15]. Erythrocyte membrane damage mechanics associated with the phosphorylation during merozoite invasion may be predicted and analysed *in silico*, e.g. with analytical models or finite element simulations. However, such *in silico* studies require the constitutive models of the biological material involved.

Therefore, this study aims to develop a constitutive model for the remodelling of the erythrocyte membrane to enable mechanistic *in silico* investigations of the merozoite invasion into erythrocytes. Continuum damage mechanics is used to represent the phosphorylation of spectrin, which is thought to alter the protein conformation, influence the dissociation of proteins, cause dissociation of protein skeleton from lipid bilayer, and induce a membrane bending moment.

## 2 Methods and materials

### 2.1 Mathematical description of the constitutive behaviour of the erythrocyte membrane damaged by phosphorylation

The hyperelastic constitutive response of a healthy erythrocyte membrane is defined by the Helmholtz free-energy function Ψ_h_ [16], a thermodynamic potential that measures the work obtainable from an isochoric and isothermal thermodynamic system [17]:

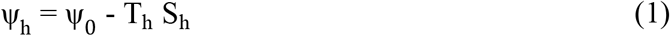

T_h_ is the absolute temperature of the surrounding modelled as a heat bath, S_h_ is the entropy of the system. and Ψ_0_ is the system’s internal energy represented using the Mooney Rivlin strain energy density function per reference volume of the intact erythrocyte membrane,

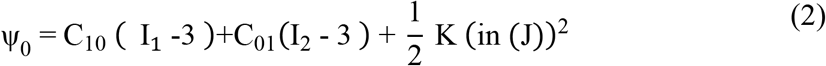

where C_10_ and C_01_ are material parameters, K is the compressibility parameter, J is the total volume ratio, I_1_ and I_2_ are the first and second deviatoric strain invariant of the left Cauchy-Green deformation tensor.

During malaria merozoite invasion, the erythrocyte membrane is deformed mainly due to the mechanical loads, exerted by the merozoite’s actomyosin machinery and external sources such as blood pressure, whereas heat transfer is negligible. Hence the energetics associated with entropy was not considered, and the Helmholtz free energy function for the erythrocyte membrane deformation was only represented as internal energy.

The damage induced by the merozoite was modelled by modifying the strain energy density function of the erythrocyte membrane. Since the erythrocyte membrane comprises primarily an elastic spectrin network and can be considered an elastic, isotropic, and nearly incompressible continuum, the strain energy density function is usually presented in a decoupled form comprising deviatoric and isochoric terms [18].

The combination of incompressibility and large deformation of a nearly incompressible hyperelastic material presents difficulties for a displacement-based finite element method as the constraint J = det F = 1 on the deformation field is highly nonlinear [19]. To overcome this challenge, a displacement-based finite element scheme must invoke a small change measure of volumetric deformation. Consequentially, the deformation gradient was decomposed into the dilatational and deviatoric parts to apply separate numerical treatments to each part [19].

Therefore, the deformation gradient **F** and the left Cauchy-Green strain tensor **B** were divided into the volume-changing (dilatational) and the volume-preserving (distortional) parts, an approach often used in elasto-plasticity [20]. The strain energy density function of the isotropic erythrocyte membrane was expressed in terms of the left Cauchy deformation tensor as:

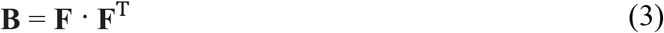

With **F** = **R U** and **F**^T^ = **R**^T^ **U**^T^, where **R** is a rotation matrix, and **U** is a stretch tensor, Eqn. (3) becomes:

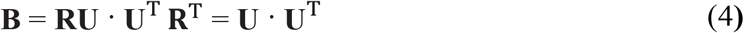

showing that the left Cauchy deformation tensor **B** is a stretch tensor and isotropic.

Hence the strain energy density function Ψ of a damaged erythrocyte membrane can be written in terms of invariants of the left Cauchy-Green deformation tensor:

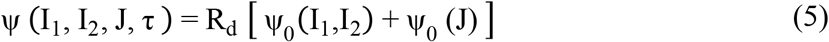

with

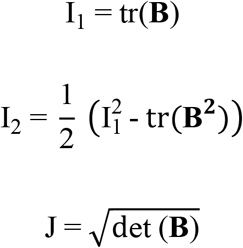

where **B** is the left Cauchy-Green tensor, R_d_ is the damage parameter, Ψ_0_ is the strain energy density of an intact erythrocyte membrane, and τ is the indentation time in seconds.

A continuum damage mechanics framework for material stiffness deterioration suitable for implementation in Abaqus Explicit was used to simulate chemical damage induced by a merozoite during the entry process. The model borrows concepts from various strain-based damage models for soft biological materials and biodegradable polymers where constitutive hydrolytic degradation and time-dependent behaviour are described [21].

The phosphorylation of the critical membrane skeleton proteins was assumed to induce alterations to both the deviatoric and the volumetric components of the strain energy density function. As a result, the elastic modulus and compressibility of the erythrocyte membrane are altered. The shear modulus μ and compressibility K of the erythrocyte membrane are:

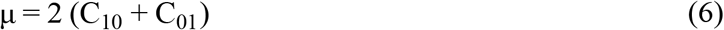

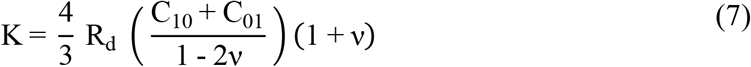

where R_d_ is a damage function, and ν is the Poisson’s ratio. R_d_ is a function of chemical and mechanical damage parameters and is defined as:

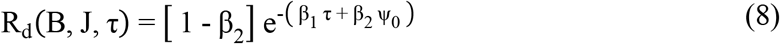

where B and J denote the left Cauchy green tensor and the volumetric strain of the intact erythrocyte membrane, respectively, τ is the simulation time, and Ψ_0_ is the strain energy density per reference volume of the intact erythrocyte membrane, β_1_ is the chemical damage parameter, and β_2_ is the mechanical damage parameter.

For β_2_ = 0, the damage mode is purely chemical, i.e., damage to the erythrocyte membrane is not due to deformation. Thus, β_1_ represents chemical damage due to various modes of phosphorylation, and β_2_ represents mechanical damage associated with the tearing or rupturing of the protein chains in the erythrocyte membrane skeleton. The erythrocyte’s membrane near incompressibility was defined with a Poisson’s ratio ν = 0.499. For incompressible materials, the contribution of the volumetric strain energy density function is neglected since J = 1. Since the erythrocyte membrane is nearly incompressible, the deviatoric strain invariants are:

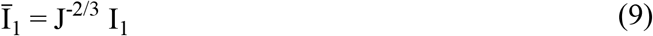

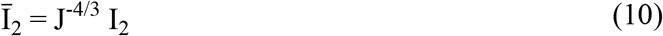

The variation of the strain energy potential is, by definition, equal to the internal virtual work per reference volume V_0,_ and this can be written as:

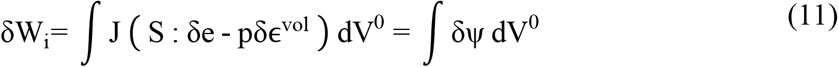

where

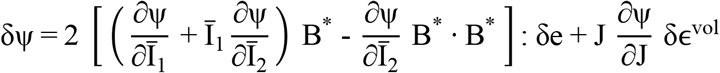

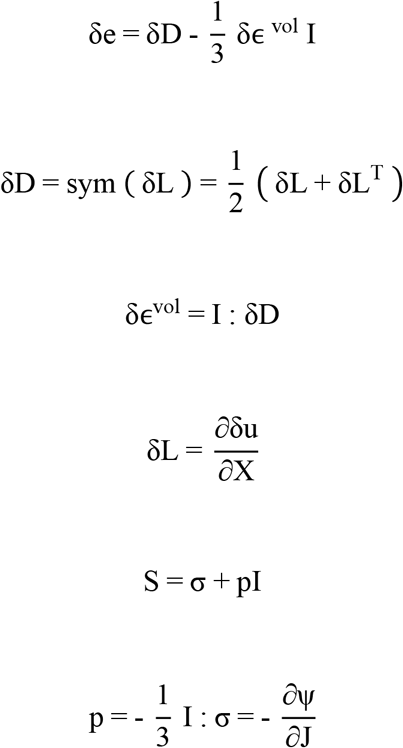

Hence the deviatoric stress can be rewritten as:

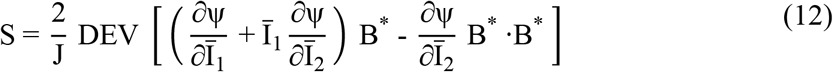

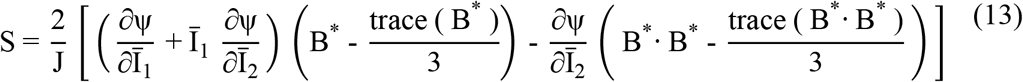

Therefore, the stress (deviatoric stress and volumetric stress) can be written as

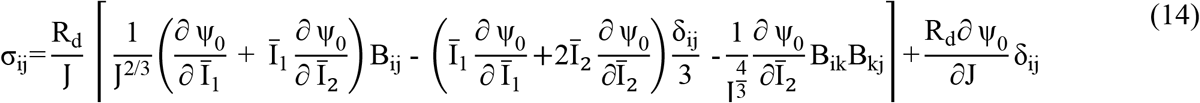

and further as

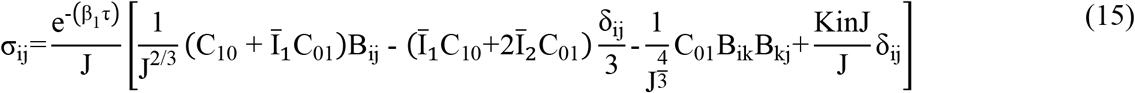

where δ is the Kronecker delta function. Ψ_0_ is the Mooney Rivlin strain energy density function per unit reference volume of the intact erythrocyte membrane:

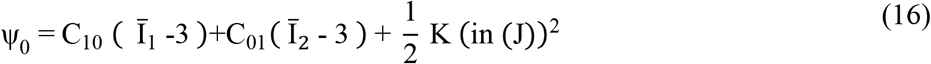

### 2.2 Implementation and verification of the erythrocyte membrane damage model in a finite element code

The erythrocyte membrane damage model was implemented as a VUMAT subroutine and used with a single shell element model to simulate merozoite-induced damage. The erythrocyte membrane damage model VUMAT subroutine is available as supplementary data.

The implementation starts with verifying the accuracy of the developed VUMAT subroutine by comparing the true stress obtained with the VUMAT subroutine and the built-in Mooney Rivlin law for an intact erythrocyte membrane with the chemical damage parameter β_1_ = 0. For this case, the constitutive responses of the developed VUMAT subroutine and the built-in Mooney Rivlin law are expected to be the same. For the verification, a single shell element was subjected to an equi-biaxial strain of 1.1, see Figure 1.

**Figure 1:**
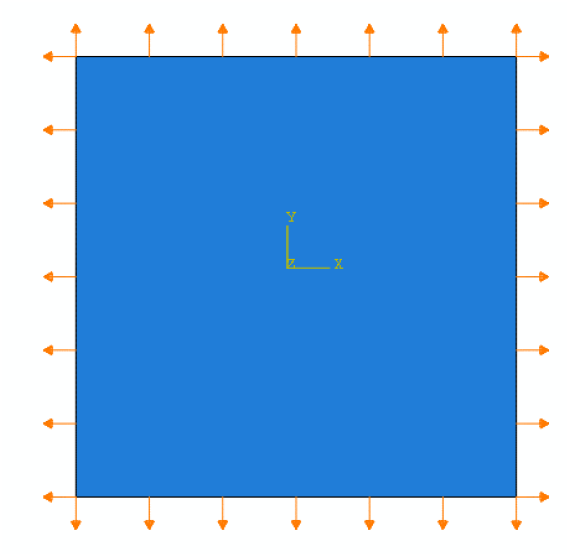
Biaxially loaded shell element used to verify the VUMAT subroutine.

After verifying the developed erythrocyte membrane damage model for β_1_ = 0, it was used in the single shell element model to simulate various degrees of chemical damage to evaluate the stability of the developed erythrocyte membrane damage model using Drucker’s stability criterion.

### 2.3 Representing erythrocyte membrane damage in alternative constitutive models

In some instances, implementing the erythrocyte membrane damage model may not be desired or possible. The suitability of other Abaqus built-in constitutive models to represent erythrocyte membrane damage was determined for such cases.

An equibiaxial tensile test was simulated in the single element model in Abaqus with the Evaluate option, which used nominal stress and strain data obtained from the developed erythrocyte membrane damage model for β_1_ = 0, 0.49, 1 and 2.7.

The following constitutive models were considered.

Ogden model:

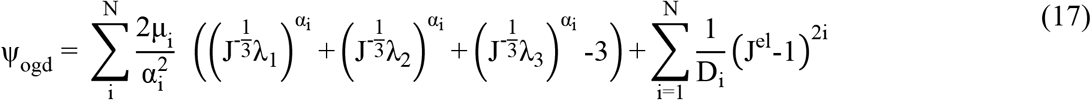

where Ψ_ogd_ is the strain energy per reference volume, λ_i_ (i = 1, 2, 3) are the principal stretches, N is the number of material parameters, μ_i_, α_i_, D_i_ are temperature-dependent material parameters, and J^el^ is the elastic volume ratio.

Yeoh model:

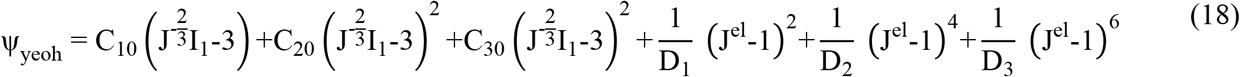

with 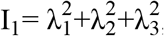, where Ψ_yeoh_ is the strain energy per reference volume, C_i0_ and D_i_ are temperature-dependent material parameters, and λ_i_ (i = 1, 2, 3) are the principal stretches.

Reduced Polynomial model:

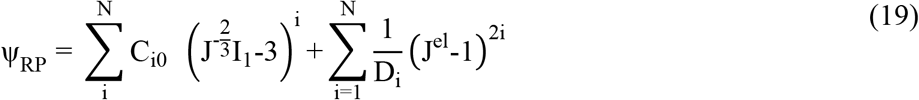

where Ψ_RP_ is the strain energy per reference volume, N is the number of material parameters, C_i0_ and D_i_ are temperature-dependent material parameters, and I_1_ is the first strain invariant.

## 3 Results

### 3.1 Erythrocyte membrane damage model

The true stresses defined by the Abaqus built-in Mooney Rivlin model and the erythrocyte membrane damage model with the parameter values provided in Table 1 agreed well for the intact membrane, i.e. for β_1_ = 0, and for true strain between 0 and 1.1 (see Figure 2 a). This agreement indicates that the developed VUMAT subroutine was accurately implemented.

**Table 1:** Material parameter values to represent an intact erythrocyte membrane with the erythrocyte membrane damage model

| $D_1$ (mm<sup>2</sup>/N) | $C_{10}$ (μN/mm<sup>2</sup>) | $C_{01}$ (μN/mm<sup>2</sup>) | $\beta_1$ |
| --- | --- | --- | --- |
| 12 | 152 | 15.2 | 0 |

**Figure 2:**
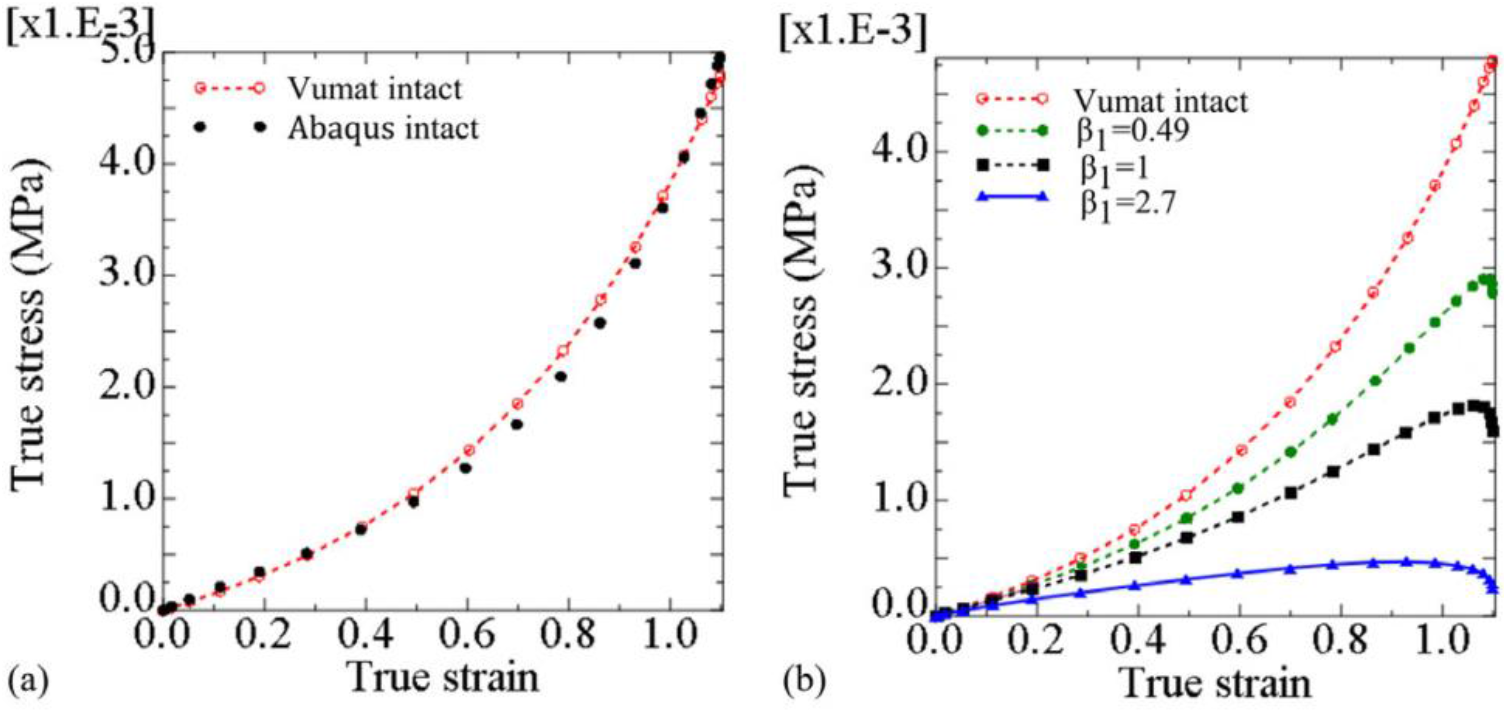
(a) True stress versus true strain in a shell element determined with the Abaqus built-in Mooney Rivlin model (‘Abaqus intact’) and the erythrocyte membrane damage model (‘Vumat intact’) for β_1_ = 0, and (b) true stress versus true strain determined with the erythrocyte membrane damage model showing the stress decrease with initiation and increase of erythrocyte membrane damage from β_1_ = 0 to 2.7.

When the damage parameter was changed from the intact state with β_1_ = 0 to the damaged state with β_1_ = 2.7, the in-plane true stress decreased for any given true strain (Figure 2 b). For a true strain of 1.1, the in-plane true stress decreased from 0.0048 MPa for β_1_ = 0 to 0.0028 MPa for β_1_ = 0.49, 0.0016 MPa for β_1_ = 1, and 0.00025 MPa for β_1_ = 2.7.

### 3.2 Representation of erythrocyte membrane damage with Ogden, Yeoh, and Reduced Polynomial constitutive models

The erythrocyte membrane stress predicted with the Ogden, Yeoh, and Reduced Polynomial models agreed well with the developed erythrocyte membrane damage model for the intact and damaged states with β_1_ = 0, 0.49, 1, and 2.7 (Figure 3 a to d). The parameter values for the Ogden, Yeoh, and Reduced Polynomial model for the different membrane states are provided in the Appendix, Table A.1 to Table A.12.

**Figure 3:**
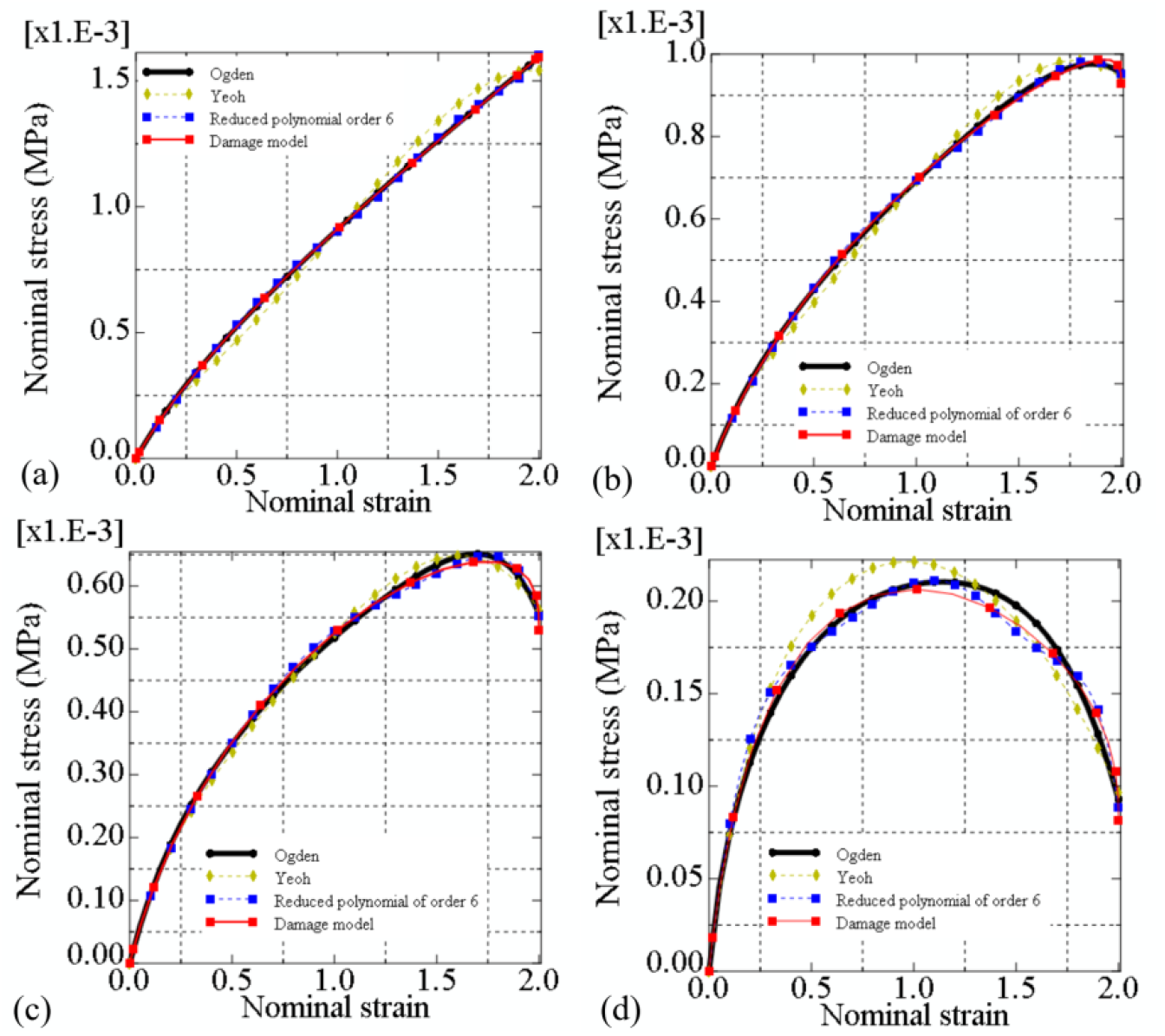
Erythrocyte membrane damage represented by the Ogden, Yeoh, and Reduced polynomial materials models at various damage states in a single shell element, for β_1_ = 0 (a), β_1_ = 0.49 (b), β_1_ = 1 (c), and β_1_ = 2.7 (d).

The Abaqus material evaluation indicated that for the intact erythrocyte membrane (β_1_ = 0), the Ogden model is the least stable of the three evaluated models. The Ogden model is unstable in biaxial tension when the biaxial nominal strain exceeds 1.05. In contrast, the Yeoh strain energy function, Reduced Polynomial strain energy function, and developed erythrocyte membrane damage model are stable in biaxial tension for the entire range of the biaxial nominal strain up to 2.0 (Figure 3 a).

For β_1_ = 0.49, the Ogden model is the least stable as it becomes unstable for a biaxial nominal strain greater than 1.75. The Yeoh and the Reduced Polynomial models become unstable when the biaxial nominal strain exceeds 1.96 and 1.98, respectively (Figure 3 b).

For β_1_ = 1, the Ogden model has the lowest stability threshold with a nominal biaxial strain of 1.74 (Figure 3 c). The stability threshold of the Yeoh and Reduced Polynomial models are at biaxial strains of 1.8 and 1.85, respectively.

For the most severe erythrocyte membrane damage with β_1_ = 2.7, the erythrocyte membrane damage model has the lowest stability threshold of a nominal strain of 1.19 (Figure 3 d and Table 2). The Ogden, Yeoh, and Reduced Polynomial models become unstable when the biaxial nominal strain exceeds 1.5, 1.36, and 1.34, respectively. The strain stability threshold of all four models is substantially lower for β_1_ = 2.7 than for β_1 =_ 0.49 and 1, indicating poor stability at extensive erythrocyte membrane damage.

**Table 2:** Strain stability threshold of various constitutive models for different erythrocyte membrane damage states β_1_. (EMD: Erythrocyte membrane damage)

| Material model | Stability threshold |  |  |  |
| --- | --- | --- | --- | --- |
| | $\beta_1 = 0$ | $\beta_1 = 0.49$ | $\beta_1 = 1$ | $\beta_1 = 2.7$ |
| EMD | stable | 1.98 | 1.89 | 1.19 |
| Ogden | 1.05 | 1.75 | 1.74 | 1.5 |
| Yeoh | stable | 1.96 | 1.8 | 1.36 |
| Reduced Polynomial | stable | 1.98 | 1.85 | 1.34 |

## 4 Discussion

To date, experimental investigations related to erythrocytes’ invasion have predominantly focused on the role of parasite adhesins, signalling pathways, and the identity of binding receptors on the erythrocyte surface. Erythrocyte membrane damage mechanics associated with the invasion process has received limited attention [22].

The erythrocyte membrane damage model developed in the current study utilises the hyperelastic Mooney Rivlin constitutive law, complemented with an exponential damage function to account for membrane remodelling due to phosphorylation. Despite limited knowledge of merozoite-induced damage, the model allowed the representation of various amounts of damage.

One constraint of hyperelastic constitutive models is their instability when strain is inversely related to stress. One way to assess a model’s stability is using Drucker’s criterion. However, this method has challenges since some hyperelastic models can be Drucker-unstable for small and large strains when subjected to different loading conditions. Hence, common practice is using a model with a known stability threshold or validity range. For example, the Mooney Rivlin model has a known validity range for equi-biaxial logarithmic strain up to 138% [23].

The stability threshold of the developed constitutive damage model was analytically determined using Drucker’s stability criteria and had similar stability thresholds as the Yeoh and Reduced Polynomial models. The initiation and increase of damage by changing β_1_ from 0 to 2.7 decrease the nominal strain stability threshold of the developed erythrocyte damage model from a stable state to 1.19. The decrease in the maximum stress represents stiffness reduction when damage is induced (Figure 2 b), indicating that the developed damage model accurately mimics the phosphorylation of the key erythrocyte membrane skeleton.

The Ogden, Yeoh, and Reduced Polynomial constitutive models could represent the constitutive response of the developed erythrocyte membrane damage model for β_1_ ranging from 0 to 2.7. Thus, these models can simulate various degrees of material damage to the erythrocyte membrane if implementing the new erythrocyte membrane model is not feasible.

The main limitation of the developed erythrocyte membrane damage model is the possible instability of the underlying Mooney Rivlin law at low strain. Furthermore, the second invariant of the left Cauchy tensor of the Mooney Rivlin law can make the developed model unstable in certain loading conditions. However, the decrease of the erythrocyte membrane stability threshold predicted with the developed model for increasing damage amount corresponds with reports that the phosphorylation of key protein elements of the erythrocyte skeleton leads to an unstable erythrocyte membrane.

## 5 Conclusions

The developed model can represent different degrees of damage to the erythrocyte membrane through phosphorylation induced by a malaria merozoite. The erythrocyte membrane damage model will enable mechanistic *in silico* investigations of the invasiveness of malaria merozoites that are, for example, useful in antimalaria drug development and screening. With suitable parameterisation of the damage variable, the developed damage model can elucidate the energetics of erythrocyte membrane damage which is challenging to investigate experimentally. The model also offers the potential for extension to study chemical and strain-based damage processes and repair mechanisms of other biological cells.

### Funding

This research was supported financially by the National Research Foundation, South Africa (grants CPRR14071676206 and IFR14011761118 to TF) and the South African Medical Research Council (grant SIR328148 to TF), and grants from the World Bank to the University of Malawi. The funders had no role in study design, data collection and analysis, the decision to publish, or the preparation of the manuscript. Any opinions, findings, conclusions, or recommendations expressed in this publication are those of the authors and do not necessarily represent the official views of the funding agencies.

### Data availability

Software used and data supporting the results presented in this article are available on the University of Cape Town’s institutional data repository (ZivaHub) under https://doi.org/10.25375/uct.22722346 as Msosa C, Abdalrahman T, Franz T. Software code and data for “A constitutive model for the remodelling erythrocyte membrane skeleton during the active invasion by the malaria merozoite”, Cape Town, ZivaHub, 2023, DOI 10.25375/uct.22722346.

## Nomenclature

B: Left Cauchy deformation tensor
C_01_: Material parameter
C_01_: Material parameter
E: Elastic modulus
F: Deformation gradient tensor
I_1_: First strain invariant
I_2_: Second strain invariant
I_i_^J^: Internal force vector
K: Compressibility of the erythrocyte membrane
R: Rotation matrix
R_d_: Damage function
S_h_: Entropy of a system
T_h_: Absolute temperature of a system
U: Stretch tensor
U_h_: Internal energy of a system
β_1_: Chemical damage parameter
β_2_: Strain or mechanical damage parameter
N: The number of material parameters for the Ogden model
δ: Kronecker delta function
μ: Shear modulus
ν: Poisson’s ratio
ψ_0_: Strain energy density of intact erythrocyte membrane
ψ_RP_: Strain energy density of the Reduced polynomial model
ψ_yeoh_: Strain energy density of the Yeoh model
ψ_ogd_: Strain energy density of the Ogden model
ψ_h_: Helmholtz free energy function
μ_i_: Temperature-dependent material parameter,
α_i_: Temperature-dependent material parameter
D_i_: Temperature-dependent material parameter
J^el^: Elastic volume ratio

## Appendix

The tables below present the material parameter values to represent a erythrocyte membrane with the Ogden, Yeoh, and Reduced Polynomial model for no membrane damage β_1_, = 0 (Tables A.1 to A.3) and membrane damage with β_1_ = 0.49 (Tables A.4 to A.6), β_1_ = 1.0 (Tables A.7 to A.9), and β_1_ = 2.7 (Tables A.10 to A.12).

**Table A.1:**
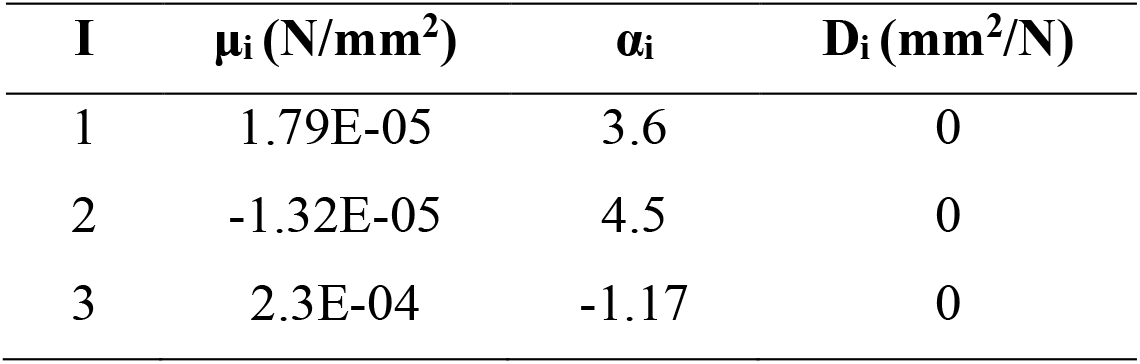
Material parameters for the Ogden strain energy function of order three, evaluated from the developed damage model nominal stress-strain data when β_1_ = 0.

| **I** | **$\mu_i$ (N/mm<sup>2</sup>)** | **$\alpha_i$** | **$D_i$ (mm<sup>2</sup>/N)** |
| --- | --- | --- | --- |
| 1 | 1.79E-05 | 3.6 | 0 |
| 2 | -1.32E-05 | 4.5 | 0 |
| 3 | 2.3E-04 | -1.17 | 0 |

**Table A.2:** Material parameters for the Yeoh strain energy function of order three, evaluated from the developed damage model when β_1_ = 0.

| **i** | **$D_i$ (mm<sup>2</sup>/N)** | **$C_{10}$ (N/mm<sup>2</sup>)** | **$C_{20}$ (N / mm<sup>2</sup>)** | **$C_{30}$ (N/mm<sup>2</sup>)** |
| --- | --- | --- | --- | --- |
| 1 | 0 | 1.32E-04 |  |  |
| 2 | 0 |  | 1.25E-05 |  |
| 3 | 0 |  |  | -3.7E-07 |

**Table A.3:** Material parameters for the Reduced polynomial strain energy function of order six, evaluated from the developed damage model when β_1_ = 0. Additional material parameters are D_1_ = D_2_ = D_3_ = D_4_ = D_5_ = D_6_ = 0.

| **$C_{10}$ (N/mm<sup>2</sup>)** | **$C_{20}$ (N/mm<sup>2</sup>)** | **$C_{30}$ (N/mm<sup>2</sup>)** | **$C_{40}$ (N/mm<sup>2</sup>)** | **$C_{50}$ (N/mm<sup>2</sup>)** | **$C_{60}$ (N/mm<sup>2</sup>)** |
| --- | --- | --- | --- | --- | --- |
| 1.24E-04 | 3.23E-05 | -5.8E-06 | 5.9E-07 | -3E-08 | 5.95E-10 |

**Table A.4:** Material parameters for the Ogden strain energy function of order three, evaluated from the developed damage model when β_1_ = 0.49.

| **i** | **$\mu_i$ (N/mm<sup>2</sup>)** | **$\alpha_i$** | **$D_i$ (mm<sup>2</sup>/N)** |
| --- | --- | --- | --- |
| 1 | 5.1E-05 | 2.87 | 0 |
| 2 | -1.4E-09 | 13.4 | 0 |
| 3 | 1.7E-04 | -0.88 | 0 |

**Table A.5:**
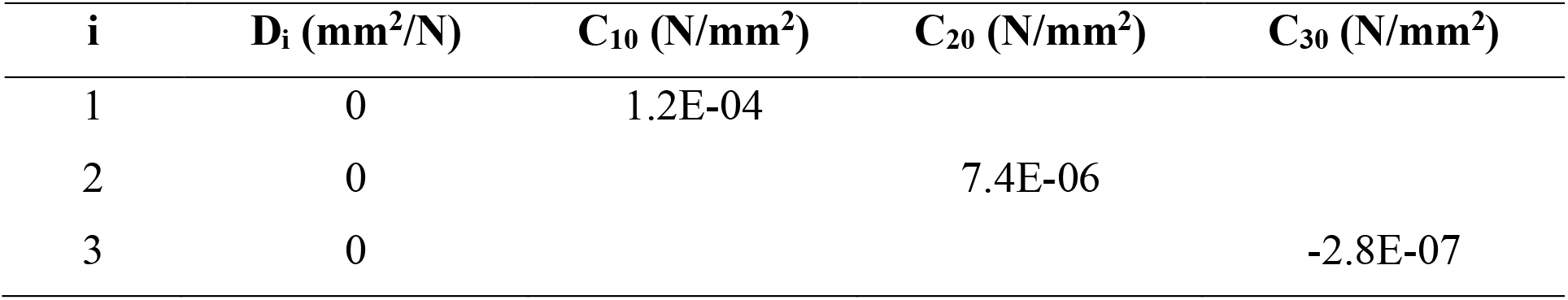
Material parameters for the Yeoh strain energy function of order three, evaluated from the developed damage model when β_1_ = 0.49.

| **i** | **D<sub>i</sub> (mm<sup>2</sup>/N)** | **C<sub>10</sub> (N/mm<sup>2</sup>)** | **C<sub>20</sub> (N/mm<sup>2</sup>)** | **C<sub>30</sub> (N/mm<sup>2</sup>)** |
| --- | --- | --- | --- | --- |
| 1 | 0 | 1.2E-04 |  |  |
| 2 | 0 |  | 7.4E-06 |  |
| 3 | 0 |  |  | -2.8E-07 |

**Table A.6:** Material parameters for the reduced polynomial strain energy function of order six, evaluated from the developed damage model when β_1_ = 0.49. Additional material parameters are D_1_ = D_2_ = D_3_ = D_4_ = D_5_ = D_6_ = 0.

| **C<sub>10</sub>** | **C<sub>20</sub>** | **C<sub>30</sub>** | **C<sub>40</sub>** | **C<sub>50</sub>** | **C<sub>60</sub>** |
| --- | --- | --- | --- | --- | --- |
| **(N/mm<sup>2</sup>)** | **(N/mm<sup>2</sup>)** | **(N/mm<sup>2</sup>)** | **(N/mm<sup>2</sup>)** | **(N/mm<sup>2</sup>)** | **(N/mm<sup>2</sup>)** |
| 1.17E-04 | 1.8E-05 | -2.9E-06 | 2.5E-07 | -1.1E-08 | 1.7E-10 |

**Table A.7:**
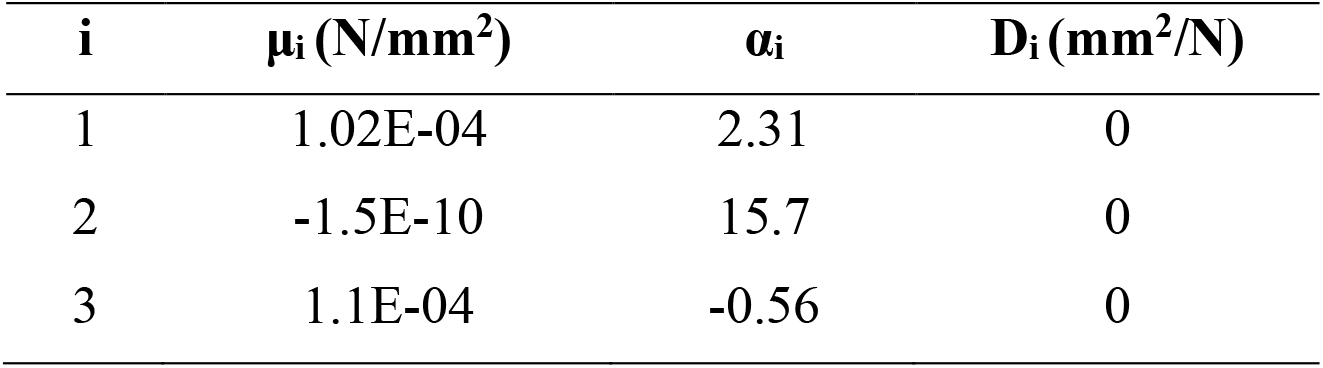
Material parameters for the Ogden strain energy function of order three, evaluated from the developed damage model when β_1_ = 1.

| **i** | **$\mu_i$ (N/mm<sup>2</sup>)** | **$\alpha_i$** | **D<sub>i</sub> (mm<sup>2</sup>/N)** |
| --- | --- | --- | --- |
| 1 | 1.02E-04 | 2.31 | 0 |
| 2 | -1.5E-10 | 15.7 | 0 |
| 3 | 1.1E-04 | -0.56 | 0 |

**Table A.8:** Material parameters for the Yeoh strain energy function of order three, evaluated from the developed damage model when β_1_ = 1.

| **i** | **D<sub>i</sub> (mm<sup>2</sup>/N)** | **C<sub>10</sub> (N/mm<sup>2</sup>)** | **C<sub>20</sub> (N/mm<sup>2</sup>)** | **C<sub>30</sub> (N/mm<sup>2</sup>)** |
| --- | --- | --- | --- | --- |
| 1 | 0 | 1.2E-04 |  |  |
| 2 | 0 |  | 3.5E-06 |  |
| 3 | 0 |  |  | -1.8E-07 |

**Table A.9:** Material parameters for the reduced polynomial strain energy function of order six, evaluated from the developed damage model when β_1_ = 1. Additional material parameters are D_1_ = D_2_ = D_3_ = D_4_ = D_5_ = D_6_ = 0.

| **C<sub>10</sub>** | **C<sub>20</sub>** | **C<sub>30</sub>** | **C<sub>40</sub>** | **C<sub>50</sub>** | **C<sub>60</sub>** |
| --- | --- | --- | --- | --- | --- |
| **(N/mm<sup>2</sup>)** | **(N/mm<sup>2</sup>)** | **(N/mm<sup>2</sup>)** | **(N/mm<sup>2</sup>)** | **(N/mm<sup>2</sup>)** | **(N/mm<sup>2</sup>)** |
| 1.1E-04 | 7.5E-06 | -9.09E-07 | 3.16E-08 | 1.03E-09 | -6.95E-11 |

**Table A.10:**
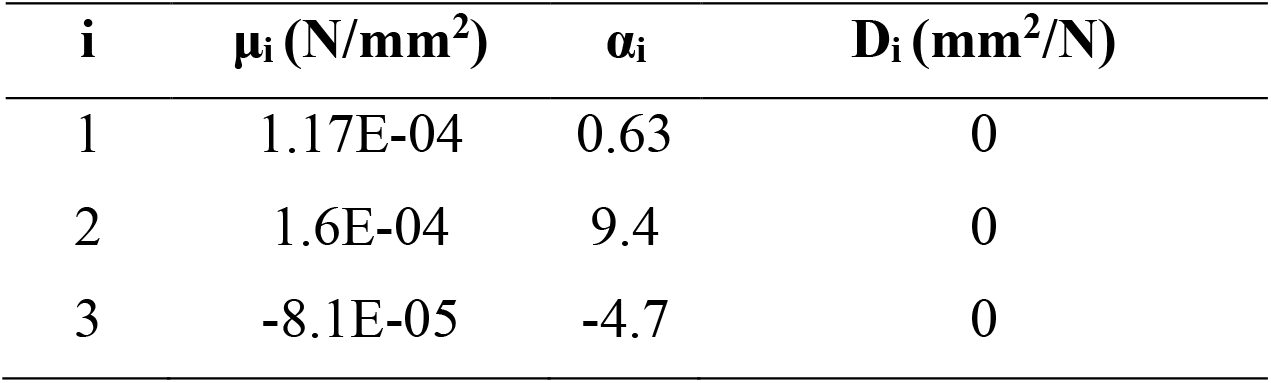
Material parameters for the Ogden strain energy function of order three, evaluated from the developed damage model when β_1_ = 2.7.

| **i** | **$\mu_i$ (N/mm<sup>2</sup>)** | **$\alpha_i$** | **$D_i$ (mm<sup>2</sup>/N)** |
| --- | --- | --- | --- |
| 1 | 1.17E-04 | 0.63 | 0 |
| 2 | 1.6E-04 | 9.4 | 0 |
| 3 | -8.1E-05 | -4.7 | 0 |

**Table A.11:**
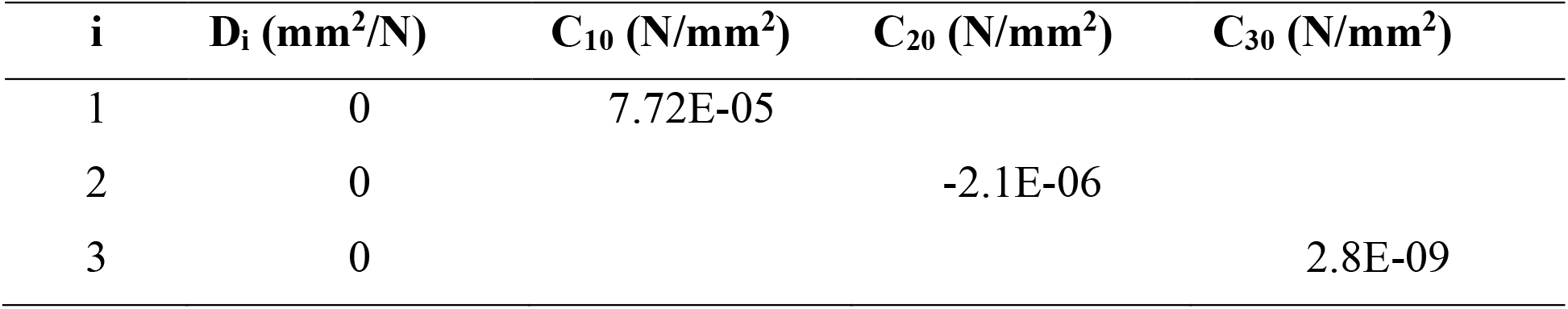
Material parameters for the Yeoh strain energy function of order three, evaluated from the developed damage model when β_1_ = 2.7.

**Table A.12:** Material parameters for the reduced polynomial strain energy function of order six, evaluated from the developed damage model when β_1_ = 2.7. Additional material parameters are D_1_ = D_2_ = D_3_ = D_4_ = D_5_ = D_6_ = 0.

| **$C_{10}$** | **$C_{20}$** | **$C_{30}$** | **$C_{40}$** | **$C_{50}$** | **$C_{60}$** |
| --- | --- | --- | --- | --- | --- |
| **(N/mm<sup>2</sup>)** | **(N/mm<sup>2</sup>)** | **(N/mm<sup>2</sup>)** | **(N/mm<sup>2</sup>)** | **(N/mm<sup>2</sup>)** | **(N/mm<sup>2</sup>)** |
| 8.5E-05 | -1.0E-05 | -1.98E-06 | -2.27E-07 | 1.25E-08 | -2.63E-10 |

